# Loss of FANCM impairs primordial germ cell differentiation *in vitro*

**DOI:** 10.64898/2026.02.09.704790

**Authors:** Amy V. Kaucher, Hannah R. Schorle, Bárbara Ribeiro Parreira, Ruiqi Sun, Kathleen R. Stewart-Morgan, Anja Pinborg, Eva R. Hoffmann, Jason A. Halliwell

## Abstract

Pathogenic variants in *FANCM* have been implicated in premature ovarian insufficiency, suggesting a critical role for FANCM in early germline development. To investigate this gene’s function in a controlled system, we evaluated whether a mouse *in vitro* primordial germ cell (PGC) model could be used to interrogate the consequences of FANCM loss on germ cell specification. Using CRISPR-Cas9 gene editing, we introduced a premature stop codon into exon 1 of *Fancm* in mouse embryonic stem cells. These edited cells were differentiated into epiblast-like cells and subsequently into primordial germ cell-like cells (PGCLCs), which were assessed for their developmental competence. After successful generation of *in vitro*-derived PGCLCs, we found that loss of FANCM markedly reduced PGCLC formation, consistent with previously reported *in vivo* phenotypes. Our findings demonstrate that this *in vitro* system provides a tractable platform for dissecting gene function in otherwise inaccessible stages of germline development. Moreover, our data suggest that FANCM is required at the earliest stages of PGC specification, potentially as early as the developmental window equivalent to embryonic day 8.5, a period not previously examined *in vivo*.

## Introduction

Fanconi anaemia (FA) is a rare, progressive disorder first described by Guido Fanconi in the 1920s and is characterized by chromosomal instability, hematopoietic failure, and increased cancer susceptibility [1,2]. At the cellular level, FA is defined by hypersensitivity to DNA interstrand crosslinking (ICL) agents such as diepoxybutane (DEB), resulting in chromosomal breakage, a key clinical diagnostic feature [3,4]. This phenotype reflects the central role of the FA pathway in the detection and resolution of ICLs, which arise frequently during DNA replication [5,6].The clinical manifestations of FA are broad, ranging from hematopoietic defects to aberrations in reproductive development and endocrine function [7]. Both male and female FA patients frequently exhibit infertility, with >50% of women developing premature ovarian insufficiency (POI) [7–9], while men are rarely fertile and commonly present with non-obstructive azoospermia (NOA) [10,11]. Variants in a range of FA genes, including *FANCM*, a DNA translocase that remodels branched DNA structures and recruits the FA core complex to sites of stalled replication forks and DNA lesions [12,13], have been linked to premature reproductive aging in the absence of other classic FA symptoms, suggesting a critical role for these proteins in reproductive longevity [14,15].

The aetiology of premature ovarian insufficiency (POI) frequently involves defects in genes responsible for maintaining genomic stability of germ cells. Several independent studies have recently reported pathogenic variants or variants of unknown significance in *FANCM* in individuals with POI (**Supplementary Table S1**) [16–24], strengthening the link between DNA repair pathways and ovarian reserve establishment and maintenance. Additionally, a recent gene-burden analysis involving POI patients of Chinese ancestry revealed three heterozygous carriers of *FANCM* variants, indicating a strong association to POI; however, specific variant-level details were not reported [23].

POI can arise not only from accelerated follicle loss postnatally, but in theory also from a reduced establishment of the primordial follicle pool during foetal development. This pool size is ultimately determined by the number and proliferative capacity of primordial germ cells (PGCs) that successfully colonize the embryonic gonad. PGCs, the embryonic precursors of sperm and oocytes, undergo a tightly regulated programme of specification, migration, and expansion that is essential for lifelong fertility [25–28]. In the mouse, PGCs are specified from the epiblast between embryonic day (E) 6.25 and E7.25 in response to BMP signalling from extra-embryonic tissues. Following specification, PGCs emerge at the base of the allantois, migrate through the hindgut and dorsal mesentery, and colonize the developing genital ridges by E10.5–E11.5, where they undergo extensive proliferation prior to sex-specific differentiation [29–31].

Disruption of FA pathway genes, including *Fancm*, have been shown to impair early germline development and compromise fertility in animal models (**Supplementary Table S2**). Specifically, *Fancm* knockout mice display reduced PGC numbers during foetal development, driven primarily by decreased proliferative capacity rather than increased apoptosis [32,33]. These findings raise the possibility that FANCM deficiency may diminish the initial germ cell reserve, providing a developmental mechanism that could contribute to POI in humans.

*In vitro* differentiation of mouse embryonic stem cells (mESC) into primordial germ cell-like cells (PGCLC) has emerged as a powerful experimental system to model germline development [34,35]. While this platform has been widely used to study germline specification pathways, its application as a tool for gene-function analysis and disease associated variant validation remains limited. Large CRISPR-based functional screens have begun to explore germ cell developmental networks [36,37] but focused studies using defined genetic perturbations, particularly those modelling disease associated genes, have remained rare [38–45].

Functional validation of genetic variants associated with reproductive aging remains a major challenge, particularly as many loci influencing age at natural menopause are thought to act during early germline development. At germline specification, PGCs are exceedingly rare, numbering ∼40 in mice [25,46] and estimated to be even fewer in humans [47,48]. Genetic disruption of key PGC-inducing factors such as *Prdm1* (BLIMP1) results in complete germ-cell loss [46,49], while perturbation of other pathways, including FA genes, leads to substantial reductions in germ-cell number in mouse models [40,50–53]. Although *in vivo* mouse studies have provided critical insights into these processes, they are costly, technically demanding, and often limited by embryonic lethality or the need for highly specialized assays to assess early germline phenotypes. Advances in *in vitro* systems, including PGCLCs and ovarian organoids [54,55], enable stage-specific interrogation of germline development outside the gonadal niche. Here, we employ the mouse PGCLC differentiation system to assess the impact of *Fancm* loss on early germline formation, with the broader aim of evaluating the utility of PGCLCs for modelling genetic variants associated with premature reproductive aging. By examining PGCLC output following targeted *Fancm* disruption, we test whether impaired germ-cell specification or expansion represents a plausible developmental mechanism linking FANCM deficiency to reduced ovarian reserve.

## Materials and methods

### Culture of mouse embryonic stem cells

A *Blimp1*-mVenus *Stella*-ECFP (BVSC) transgenic reporter mouse embryonic stem cell line was acquired from the group of Mitinori Saitou, Kyoto University, Japan [56]. Feeder-free mESCs were maintained in serum free basal media, supplemented with L-glutamine (2 mM; Invitrogen, #14190-094), Leukaemia inhibitory factor (LIF; 1 × 10^3^ IU/mL; Sigma-Aldrich, #ESG1107), GSK-3 inhibitor (CHIR99021; 3 µM; Selleckchem, #1263), MEK/ERK inhibitor (PD0325901; 1 µM; Selleckchem, #1036), and monothioglycerol (0.00126%; Sigma-Aldrich, #M6145). Cells were cultured on 6-well tissue culture plates (Corning, #CLS3516) coated with poly-L-ornithine (0.01%; Invitrogen, #P3655) and laminin (300 ng/mL; Corning, #CLS354232). Cells were collected every third day by enzymatic disaggregation (via TrypLE Express Enzyme; Invitrogen, #12604013) and passaged to fresh wells at a plating density of 2.5 × 10^4^ cells per well of 6-well tissue culture plates. All cells were maintained at 37°C in 5% CO_2_ and 95% air in a humidified Heracell Vios 160i incubator (ThermoFisher Scientific).

### Generation of Fancm^−/−^ clones

Guide RNAs (CRISPR RNAs [crRNAs]; **Supplementary Table S3**) were designed to target the first exon of *Fancm*, adjacent to an identified protospacer adjacent motif (PAM) sequence. No repair template was used; therefore, editing relied upon nonhomologous end-joining. CRISPR-Cas9 editing was conducted according to manufacturer’s instructions (Integrated DNA Technologies). Briefly, tracr:crRNA duplexes were formed by mixing equimolar amounts of Alt-R crRNA (Integrated DNA Technologies) and Alt-R tracrRNA ATTO-550 (Integrated DNA Technologies, #1975827) in IDT Duplex Buffer (30 mM HEPES, pH 7.5, 100 mM potassium acetate; Integrated DNA Technologies, #11-01-03-01), heating to 95°C for 5 min and cooling to room temperature for 5-10 min. Ribonucleoprotein (RNP) complexes were formed by combining equimolar amounts of tracr:crRNA duplexes with Cas9 nuclease (Alt-R Streptococcus pyogenes Cas9 nuclease; Integrated DNA Technologies, #1081058) and incubated at room temperature for 20 min. Single-cell suspensions of the *Blimp1*-Venus *Stella*-ECFP transgenic reporter mouse embryonic stem cell line were transfected with a cocktail of gRNA, Cas9 enzyme, and purified ATTO protein with or without HDR template.

Cells were transfected using the Neon Transfection System (Invitrogen, MPK1096) following manufacturer’s instructions. First, cells were resuspended in Resuspension Buffer R at 1 × 10^7^ cells per mL. Electroporation Enhancer (100 μM) and 1 μL of RNP complex (described above) were combined with 10 μL of cell suspension in Resuspension Buffer. This suspension was loaded into a Neon pipette (Invitrogen, #MPP100), inserted into the electroporation tube, and electroporated for 3 cycles at 1400 V for 10 ms. Cells were immediately transferred into a Costar 12-well plate (Corning, #3513) coated with poly-L-ornithine (0.01%; Invitrogen, #P3655) and laminin (300 ng/mL; Corning, #CLS354232) and cultured overnight. The following day, transfected cells were assessed for the presence of ATTO-550 (*i.e.* proof of successful transfection) and sorted as a single cell per culture well of Costar 96-well plates (Corning, #3595) using a Sony SH800Z Cell Sorter (Sony Biotechnology). Sorted cells were maintained under mESC conditions to allow for clonal expansion.

### Bulk shallow genomic sequencing for karyotyping

Genomic DNA was isolated from potential *Fancm* CRISPR-knockout clones via DNeasy Blood and Tissue extraction kit (Qiagen, #69582). Whole genome sequencing libraries were prepared using Nextera XT DNA Library Preparation Kit (Illumina; FC-131-1096) and analysed on a NextSeq 2000 Sequencing Systems (Illumina). Downstream analyses were completed with R (4.2.1) via an in-house pipeline [57]. Sequencing reads (> 1 million, paired-end) were aligned to the mm39 genome using Rbowtie2 (version 2.3.1) [58] to create SAM files and converted into BAM files with Rsamtools (version 2.13.4) [59]. Read counts were binned along the genome (200 kb windows) using the QDNASeq package (version 1.33.1) [60] and then GC and mappability corrected. Bin counts were also normalised internally within each sample (QDNAseq [version 1.33.1] normalizeBins) and smoothed (QDNAseq [version 1.33.1] SmoothOutlierBins) for downstream copy number variation plots using ggplot2 (version 3.3.6) [61]. Chromosome abnormalities such as aneuploidy and partial chromosome copy number variants (≥5 Mbps) were visualised and reported without knowledge about the corresponding genotype result. DNA derived from epididymal mouse (C57bl/RJj; Janvier Labs) sperm was used as a quality control.

### Sanger sequencing

Genomic DNA was isolated from potential *Fancm* CRISPR-knockout clones via DNeasy Blood and Tissue extraction kit (Qiagen, #69504). Genomic DNA was amplified using Q5-Hot Start polymerase (New England Biolabs, #M0494S) from a 1,173 bp region of the first exon of the mouse *Fancm* gene spanning the gRNA interaction site. Primers sequences for the first exon region of *Fancm* can be found in **Supplementary Table S3**. The resulting PCR product was purified using QIAquick PCR Purification kit (Qiagen, #28104) and sequenced by Eurofins Genomics Europe. Generated sequences were analysed with SnapGene (Dotmatics) software, as wells as Clustal Omega [62] and ExPASy – Translate [63] tools.

### Western blotting and analysis

Protein lysates were isolated from cultured cells by using radioimmunoprecipitation assay buffer (RIPA; Invitrogen, #R0278) with cOmplete Protease Inhibitor Cocktail (1x; Roche, #04693116001), PhosSTOP Phosphatase Inhibitor Cocktail (1x; Roche, #04906837001), and benzonase (1:1000; Sigma-Aldrich, #E1014). Cells were lysed at a concentration of 2 × 10^6^ cell per mL of lysis buffer, and samples were incubated with NuPAGE 1x LDS sample loading buffer (Invitrogen, #NP0008) at 95°C for 5 min. Protein lysates in loading buffer (100 µl) were loaded onto Novex 3-8% tris-acetate gels (Invitrogen, #EA03785BOX), and gel electrophoresis was conducted using an XCell SureLock Mini-Cell electrophoresis system (Invitrogen) at 75 V for 2 h 30 min. Proteins were transferred to PVDF membranes (iBlot™ 2 Transfer Stacks, Invitrogen, #IB24002) using the iBlot 2 system (Invitrogen, #IB21001) at 25 V for 5 min. Blots were blocked for nonspecific antibody binding by incubation with 5% non-fat dry milk in Tris-buffered saline with 0.1% Tween-20 (TBS-T) for 2 h at room temperature with gentle shaking. Blots were probed with rabbit-anti human FANCM primary antibody (1:1000 dilution; Bethyl Laboratories, #A302-637A) overnight at 4°C with gentle shaking followed by washing (3x) with TBS-T. The blots were incubated with secondary goat anti-rabbit IgG antibody conjugated to horseradish peroxidase (HRP; 1:5000 dilution; Sigma Aldrich, #AP307P) for 2 h at room temperature with gentle shaking, followed by washing (3x) in TBS-T. HRP-activity was detected using Amersham ECL Select Western blotting detection reagent (Cytiva Life Sciences, #RPN2236). Digital images were captured after 30-60 s of exposure by using a GelDoc XRS+ Imaging System (Bio-Rad Laboratories). For normalization, blots were then stripped of antibodies by using Restore Western blot stripping buffer (Thermo Scientific, #21059) for 15 m and were again blocked against nonspecific antibody binding. Subsequently blots were probed with rat anti-human Tubulin alpha-1A chain (1:1000 dilution; Novus Biologicals, #NB100-1639) primary antibody overnight at 4°C and with secondary goat anti-rat IgG antibody conjugated to HRP (1:5000; Sigma Aldrich, #AP307P) for 2 h at room temperature. Again, HRP-activity was detected using Amersham ECL Select Western blotting detection reagent (Cytiva Life Sciences, #RPN2236), and digital images were captured after 30-60 s of exposure by using a GelDoc XRS+ Imaging System (Bio-Rad Laboratories). Densitometric analyses were conducted using ImageJ/Fiji software [64].

### Generation of mouse epiblast-like cell differentiation

*Blimp1*-Venus *Stella*-ECFP (BVSC) mESCs were differentiated over the span of 2 days into epiblast-like cells. Differentiation to EpiLCs was accomplished as previously detailed [54,55] by passaging 1 × 10^5^ *Blimp1*-Venus mESC to human plasma fibronectin (16.7µg/mL; Sigma-Aldrich, #FC010) coated 12-well tissue culture plates (Corning, #CLS3513), in N2B27 base media consisting of DMEM/F12 (Invitrogen, #31331028), N2 supplement (0.5x; Invitrogen, #17502001), Neurobasal medium (0.5x; Invitrogen, #21103049), B27 supplement (1x; Invitrogen, #12587081), Penicillin-Streptomycin (100 IU/mL and 100 µg/mL, respectively; Invitrogen, #15140122), and L-Glutamine (0.1 mM; Invitrogen, #25030081) supplemented with Activin A (20ng/mL; Peprotech, #120-14) and FGF2 (12ng/mL; Invitrogen, #13256-029). The cells were grown under these conditions for 42 h, with media being refreshed after 24 h of culture.

### Primordial germ cell-like cell differentiation

After 42 hours of culture, epiblast-like cells were collected and differentiated into primordial germ cell-like cells as previously described [54,55] by collecting 2 × 10^3^ EpiLCs at 42 h of differentiation culture. Collected EpiLCs were transitioned to Glasgow Minimum Essential Media (Invitrogen, #11710035) media supplemented with KnockOut Serum Replacement (KOSR; 15%; Invitrogen, #10828028), Penicillin-Streptomycin (100 IU/mL and 100 µg/mL, respectively; Invitrogen, #15140122), Sodium pyruvate (1 mM; Sigma-Aldrich, #P5280), L-Glutamine (0.1 mM; Invitrogen, #25030081), Nonessential Amino Acids (0.1 mM; Invitrogen, #11140050), and β-mercaptoethanol (0.1 mM; Invitrogen, #21985023), as well as cytokines BMP4 (500 ng/mL; R&D Systems, #Q53XC5), BMP8a (500 ng/mL; R&D Systems, #P34820), EGF (50 ng/mL; R&D Systems, #P01133), LIF (1 × 10^3^ IU/mL; Sigma-Aldrich, #ESG1107), and SCF (100 ng/mL; R&D Systems, #Q78ED8). The cells were grown in Costar 96-well, low-binding, round-bottom plates (Corning, #3799) coated with Lipidure reagent (0.5%; Amsbio, #AMS.52000012GB100G) for 2-6 days to support embryoid body formation.

### Microscopy, image processing, and image analysis

Light and fluorescent microscopy were performed using 4x, 10x, 20x, and 40x objectives on a CKX53 microscope, a SC50 camera, and CellSens software (Olympus). Visualization of the *Blimp1*-Venus fluorescent transgene was accomplished using the pE-300 LED Illumination System (CoolLED) with a standardized exposure time of 100 ms. Image analyses were conducted using ImageJ/Fiji software [64]. Representative fluorescent images were processed by converting to grayscale, enhancing contrast, and inverting colour to improve clarity of visual information. All processing was identically applied across all representative images using ImageJ/Fiji software [64].

### Flow cytometric analyses and fluorescent-activated cell sorting

Flow cytometric analyses were performed on a Sony SH800Z Cell Sorter (Sony Biotechnology), following previously reported gating strategies [56]. Fluorescent gating was based on Venus versus PE fluorescence to manage autofluorescence. As emissions of autofluorescence are in the Venus and PE channels, these cells will visualize in the diagonal; therefore, auto-fluorescent cells will be able to be distinguished from the low-Venus expressing cells. Flow cytometric data were analysed and visualized using FCS Express software (De Novo Software).

### Quantitative reverse transcriptase polymerase chain reaction

RNA was extracted from cells using RNeasy Micro Extraction Kit (Qiagen, #74104) according to the manufacturer’s instructions. RNA concentration and purity (as determined by 260/280-nm absorbance reading ratios) were measured using a Qubit 2.0 Fluorometer (Invitrogen). Only samples with a 260/280 nm ratio of greater than 1.75 were used for qRT-PCR analyses. For each sample, 1 µg of RNA was reverse transcribed with oligo(d)T priming by using Maxima First Strand cDNA Synthesis Kit for RT-qPCR with dsDNase (Thermo Fisher Scientific, #K1672). Reaction cocktails included approximately 2ng of cDNA, Power SYBR Green PCR master mix (Thermo Scientific, #4367659), and previously reported qPCR primer sequences (**Supplementary Table S3**) [65]. Relative gene expression levels were determined using a StepOnePlus Real-Time PCR system (Applied Biosystems). To make quantitative comparisons between samples, target gene expression was normalized to expression of the constitutively expressed *glyceraldehyde-3-phosphate dehydrogenase* (*Gapdh*) gene. Relative transcript abundance was determined using the formulae:

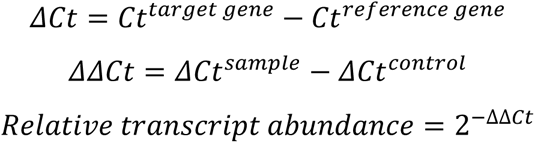

### Apoptosis assay

Apoptosis assay was performed using the eBioscience Annexin V Apoptosis Detection Kits (ThermoFisher Scientific; #88-8007-74). Cells were collected and washed with DPBS (Invitrogen, #14190169). Cells were resuspended at 5 × 10^5^ in 100 μL Binding Buffer and 5 μL of AnnexinV-APC antibody. After 10 m incubation at room temperature cells were washed in 1x Binding Buffer and resuspended in 200 μL Binding Buffer. Propidium iodide (PI; 5 μL) was added to the cell suspension and analysis was conducted using the Aurora Spectral Flow Cytometer (Cytek). Spectral data were analysed using FCS Express software (De Novo Software).

### Proliferation assays

Proliferation of mESCs was assessed by cell proliferation via manual counting. On Day 0, cells were seeded into 6-well plates (Corning, #CLS3516) at a density of 2.5 × 10^4^ cells per well and cultured via previously detailed methods. At 72 h cells were collected, resuspended in 1 mL of fresh media, and counted using a haemocytometer. Viability was assessed via exclusion of trypan blue (ThermoFisher Scientific, #15250061), and the total number of viable cells was recorded.

Proliferation of EpiLC was assessed by the same means as mESCs, with the following exceptions: EpiLCs were cultured as previously detailed herein, the initial plating concentration was 1 × 10^5^ in a 12-well culture plate (Corning, #CLS3513), and EpiLCs were collected and counted after 42 h of culture.

Proliferation of PGCLCs was analysed via CellTrace Far Red Cell Proliferation Kit (ThermoFisher Scientific, #C34564). Prior to PGCLC induction, EpiLCs were labelled for 30 m using CellTrace (1 μM). After either 48 or 96 h PGCLCs were analysed for CellTrace using the Cytek Aurora spectral flow cytometer (Cytek). Spectral data were analysed using FCS Express software (De Novo Software).

### Statistical analyses and visualization

Statistical analyses were performed using GraphPad Prism 10 (Dotmatics). Normalcy of distribution was assessed via the Shapiro-Wilk test. Levene’s test was used to evaluate equality of variances. Normally distributed data with equal variances were evaluated by unpaired t-tests with Two-stage step-up (Benjamini, Krieger, and Yekutieli) multiple comparisons correction method; ordinary one-way ANOVA with Tukey’s post-hoc test; or two-way ANOVA Dunnett’s multiple comparisons test. Normally distributed data with unequal variances were evaluated by Welch’s ANOVA with Games-Howell’s post-hoc test. Data determined to be non-normal were evaluated with Mann-Whitney or Kruskal-Wallis test with Dunnett’s post-hoc test. Effect size between two groups was calculated using Cohen’s d (*d*); whereas, between multiple groups, effect size was evaluated using Omega-squared (ω²). A P-value of ≤0.05 was considered significantly different. Visualization of data was created using GraphPad Prism 10 (Dotmatics).

### Sample size justification

The sample size was calculated based on preliminary data, using G*Power 3.1.9.7 software [66]. The parameters were defined as: effect size of 0.95, α error probability of 0.05, power of 0.8, and an allocation ratio of 1. The total number of cell lines required for this study was calculated to be 26 (13 for genotype: *Fancm*^+/+^ and *Fancm*^−/−^ knockout). However, n = 4 per genotype (n = 8 total) was chosen as the starting point. This decision took into consideration the robustness of the preliminary data. Six replicates per condition were advised for sequencing experimentation [67], but reducing the number of cell line replicates to 4 yielded an ad hoc power of 0.44. Given that this level of statistical power was deemed sufficient for the intended analysis, additional replication was considered unnecessary.

## Results

### Modelling early germline development with in vitro-derived primordial germ cells

Despite the body of evidence supporting the role of FANCM in non-syndromic POI (**Supplementary Table S1**), the mechanisms by which FANCM dysfunction contributes to ovarian failure remain unclear. We therefore sought to use the PGCLC model to characterize the functional impact of FANCM loss on cellular processes relevant to the earliest stage of germline development (*i.e*., PGCs).

To model FANCM deficiency in early germline development, an *in vitro* method of PGC specification was established [54,55]. BLIMP1, a key transcriptional regulator for PGC fate [46,49,68,69], is widely used as a germline marker for *in vitro* differentiation protocols [54,55]. Accordingly, an mESC line carrying a *Blimp1*-Venus (BV) fluorescent reporter was employed [56]. mESCs were first differentiated into epiblast-like cells (EpiLCs) under the influence of Activin A and FGF2 for two days. The resulting EpiLCs were then aggregated to form embryoid bodies in the presence of BMP4, which induces a subset of the cells to adopt a PGCLC fate over the following 6 days, while the remaining cells within the embryoid bodies differentiate into various somatic lineages (**Fig. 1a**).

**Fig. 1.**
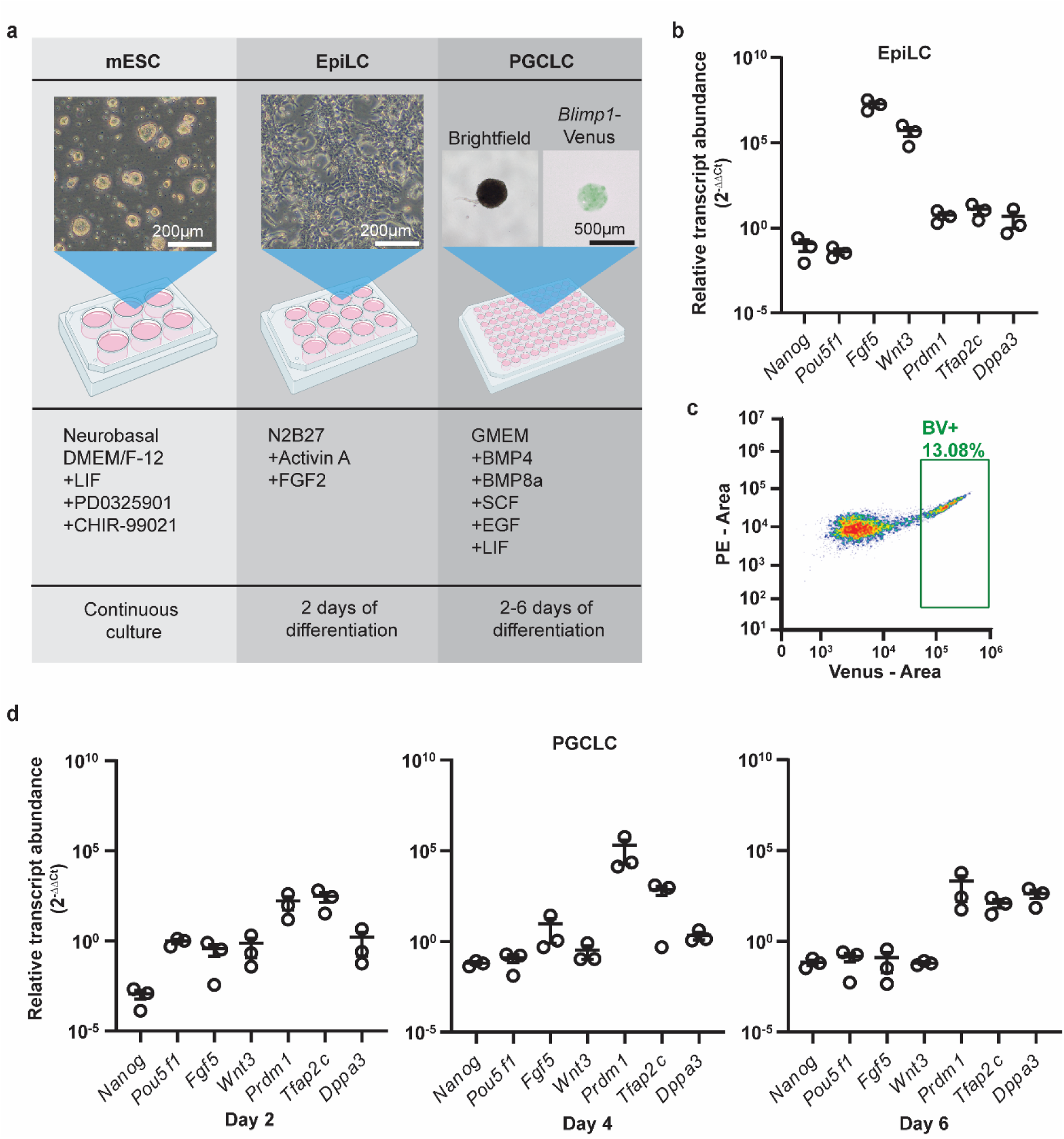
Modelling early germline development using *in vitro*-derived primordial germ cells (PGCs). (a) Workflow of mouse *in vitro* PGC-like cells (PGCLCs). Mouse embryonic stem cells (mESCs); *Blimp1*-Venus (BV); Epiblast-like cells (EpiLCs). (b) Expression of markers of pluripotent- (*i.e.*, *Nanog*, *Pou5f1*), epiblast- (*i.e.*, *Fgf5*, *Wnt3*), and germline- (*i.e.*, *Prdm1*, *Tfap2c*, and *Dppa3*) lineages of the *Blimp1*-Venus (BV) reporter line at the EpiLC stage of differentiation, normalized to the gene expression of BV mESCs. Data are presented in log2 scale. (c) Representative scatter plot and gating for flow cytometric analyses of disassociated embryoid bodies derived from BV reporter cell line on day four of PGCLC differentiation. PGCLCs are distinguished by expression of the BV transgene (BV+), indicated by rectangular gate. (d) Expression markers of pluripotent- (*i.e.*, *Nanog*, *Pou5f1*), epiblast- (*i.e.*, *Fgf5*, *Wnt3*), and germline- (*i.e.*, *Prdm1*, *Tfap2c*, and *Dppa3*) lineages of the BV+ cells at day 2, 4, and 6 of PGCLC differentiation, normalized to the gene expression of BV mESCs (log2 scale).

By day two of EpiLC differentiation, expression of pan-epiblast (*Fgf5)* and proximal epiblast (*Wnt3*) markers were upregulated, while pluripotency factors *Nanog* and *Pou5f1* were downregulated compared to mESC gene expression (**Fig. 1b**). Subsequent formation of embryoid bodies allowed further differentiation into PGCLCs over six days, and *Blimp1*-Venus positive (BV+) PGCLCs were isolated by FACS (**Fig. 1c**), and their identity was confirmed by qRT-PCR analysis, which demonstrated progressive upregulation of germline markers (*Prdm1*, *Tfap2c,* and *Dppa3)* and loss of epiblast marker expression (*Fgf5* and *Wnt3*) (**Fig. 1d**). These results confirm that the BV reporter cell line reliably generates PGCLCs under standard conditions.

### CRISPR-Cas9 mediated FANCM ablation in mESCs

To model FANCM deficiency during PGC specification *in vitro*, a CRISPR-Cas9 gene editing approach was used to introduce a premature stop codon into the first exon of *Fancm* (**Fig. 2a-c**). Western blotting with an antibody targeting residue 1-50 of FANCM identified six clones with undetectable protein, indicating successful loss-of-function editing (**Fig. 2d**; clone J, K, L, M, V, and W). Sanger sequencing confirmed the presence of a premature stop codon in four of these clones (clone J, L, M, and W), effectively ablating all of the functional domains of the protein and likely subjecting the transcript to nonsense mediated decay (**Fig. 2e, Supplementary Table S4, S5**) [70]. These clones are subsequently referred to as *Fancm*^−/−^, while four clones that were transfected with the CRISPR-Cas9 but were found to be unedited (clones E, G, I, Q) were used as *Fancm*^+/+^ controls (**Fig. 2d, e; Supplementary Table S4, S5**).

**Fig. 2.**
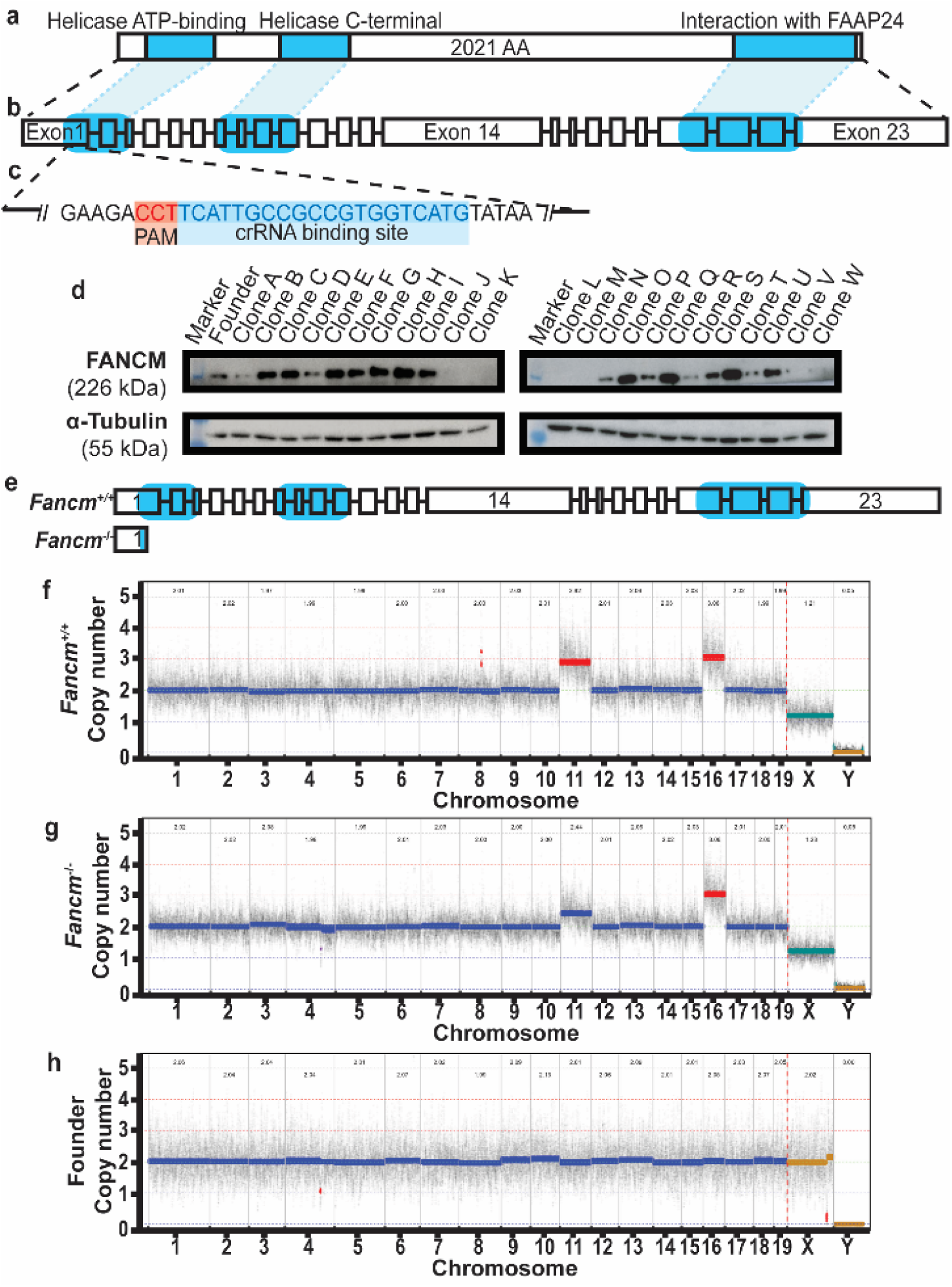
Establishing *Fancm* knock-out cell line by CRISPR-Cas9 gene editing. (a) Schematic representation of the mouse FANCM protein with annotated functional domains (highlighted, blue). (b) Structure of the full-length *Fancm* gene with coding regions corresponding to each functional domain indicated (highlighted, blue). (c) Nucleotide-level view of a portion of *Fancm* exon 1 showing PAM sequence and guide RNA (crRNA) target site for CRISPR-Cas9 genome editing. (d) Representative Western blot images of *Fancm* CRISPR-Cas9 transfected clones probed for FANCM protein (upper panel) and loading control α-Tubulin (lower panel). (e) Comparison of full-length transcripts of *Fancm*^+/+^ clones and truncated *Fancm*^−/−^ clones, the latter resulting from premature stop codon introduced in exon 1 that eliminated all functional domain coding regions. (f-h) Representative traces of whole genome chromosome copy number for BV mESCs (*Fancm*^+/+^, *Fancm*^−/−^, and founder lines, respectively) generated by low-pass whole genome sequencing.

Prolonged culture of mESC, such as during CRISPR-Cas9 genetic manipulation and clonal expansion, can select for aneuploidies that confer growth advantages *in vitro* [71,72]. To assess genomic integrity, low-pass whole genome sequencing was performed to detect aneuploidy and copy number variants. Both *Fancm*^+/+^ and *Fancm*^−/−^ clones shared identical aneuploidies affecting chromosome 11, 16 and X, which were absent in the founder line with a normal karyotype (**Figure 2f-h, Supplementary Table S6**). Although these shared aneuploidies could be a potential confounding factor, their presence in both genotypes allow attribution of any observed differences in germline specification specifically to FANCM loss, rather than to the aneuploidies.

### Loss of FANCM does not influence pluripotent or epiblast-like cell states

To examine the influence of FANCM loss on cell identity prior to PGCLC specification, mESCs and EpiLCs were first assessed. Cell morphology at the mESC (**Fig. 3a**) and EpiLC (**Fig. 3b**) stages appeared normal. Culture of *Fancm*^−/−^ mESCs for 3 days revealed no statistically significant difference in total cell numbers compared to *Fancm*^+/+^ or the euploid founder cell line (**Fig. 3c**; p = 0.1073, ordinary one-way ANOVA; ω² = 0.053, small-to-moderate effect). Similarly, after two days of differentiation, EpiLC counts did not differ between genotypes (**Fig. 3d**; p = 0.865, ordinary one-way ANOVA; ω² = 0.017, small-to-moderate effect). Annexin V and propidium iodine staining further demonstrated no significant difference in early, late, or total apoptotic fractions at either the mESC (**Fig. 3e**; p-values and effect sizes reported in **Supplementary Table S7**) or EpiLC stage (**Fig. 3f**; p-values and effect sizes reported in **Supplementary Tables S7, S8**). Gene expression analyses confirmed that pluripotency and epiblast markers were unaffected by FANCM loss (**Fig. 3g, H**; p-values and effect sizes reported in **Supplementary Table S9**). Collectively, these data indicate that both *Fancm*^+/+^ and *Fancm*^−/−^cells retain morphology, proliferation, and gene expression patterns consistent with the founder line at the pluripotent and epiblast-like stages.

**Fig. 3.**
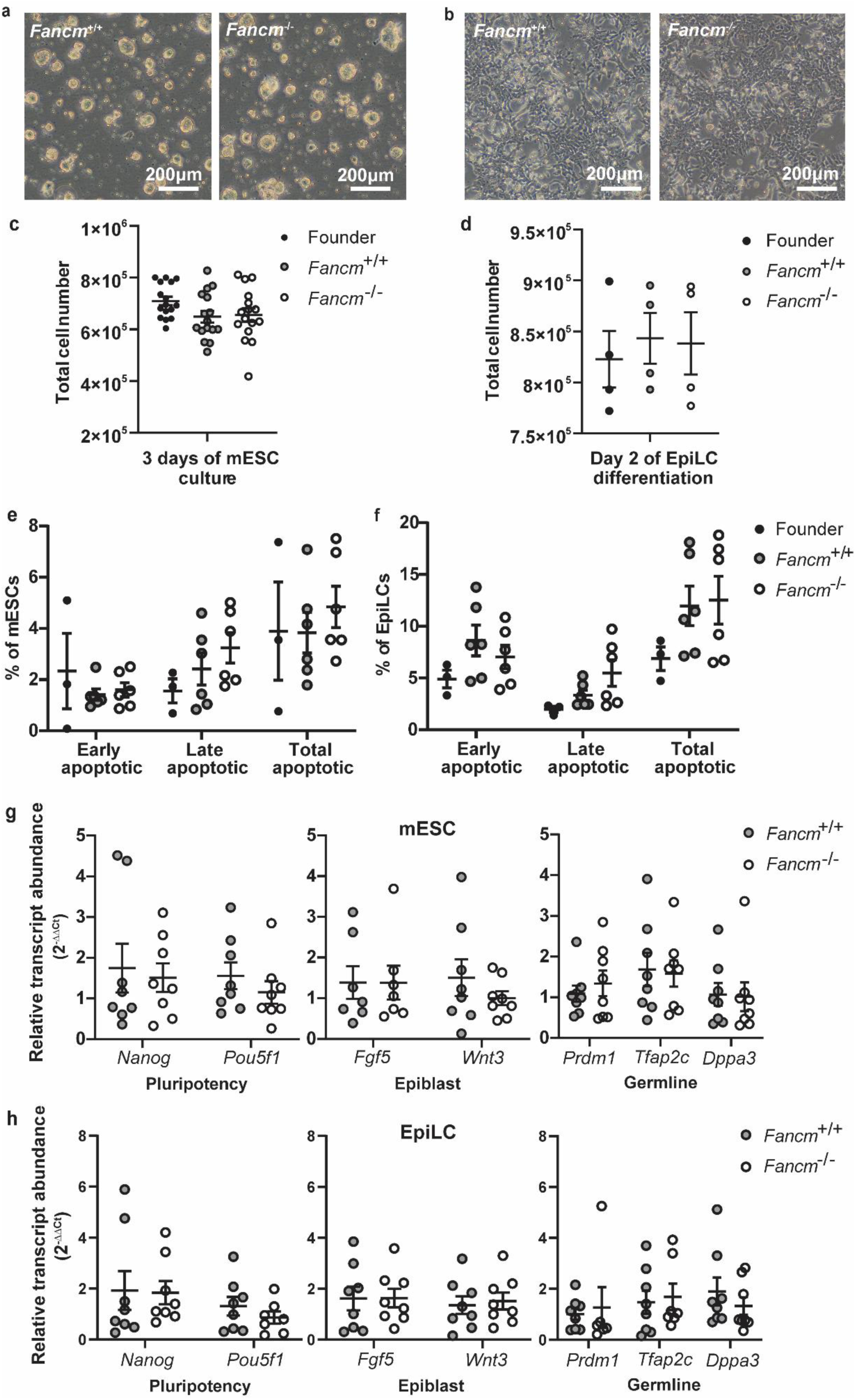
Ablating FANCM does not alter pluripotent or epiblast-like cell characteristics. (a-b) Representative brightfield images of *Fancm*^+/+^ and *Fancm*^−/−^ mESCs or EpiLCs, respectively. Scale bar = 200 µm. (c) Expansion of founder, *Fancm*^+/+^, and *Fancm*^−/−^ mESC cultures assessed by cell proliferation via manual counting over 72-hour culture period. (d) Expansion of EpiLCs measured by cell proliferation via manual counting on the second (final) day of EpiLC differentiation. (e-f) Apoptotic profile of founder, *Fancm*^+/+^, and *Fancm*^−/−^ mESCs or EpiLCs, respectively. Stages of apoptosis are defined as early (Annexin V +, PI -), late (Annexin V + and PI +) and total (Annexin V + and PI −/+) apoptotic cell fractions. (g) Relative transcript abundance of markers of pluripotent- (*i.e.*, *Nanog*, *Pou5f1*), epiblast- (*i.e.*, *Fgf5*, *Wnt3*), and germline- (*i.e.*, *Prdm1*, *Tfap2c*, and *Dppa3*) lineages of *Fancm*^+/+^ and *Fancm*^−/−^ BV reporter mESC lines, normalized to unedited founder BV mESC gene expression. Data are presented in log2 scale. (h) Relative transcript abundance of markers of pluripotent- (*i.e.*, *Nanog*, *Pou5f1*), epiblast- (*i.e.*, *Fgf5*, *Wnt3*), and germline- (*i.e.*, *Prdm1*, *Tfap2c*, and *Dppa3*) lineages of *Fancm*^+/+^ and *Fancm*^−/−^ BV reporter cells lines at the EpiLCs stage, normalized to founder EpiLC transcript abundance. Data are presented in log2 scale.

### Loss of FANCM impairs PGCLC specification

To model the impact of FANCM loss on PGC differentiation, we imaged the embryoid bodies and quantified their area from brightfield images. Aggregate size did not differ significantly among the founder, *Fancm*^+/+^, and *Fancm*^−/−^ lines at day 2 (0.089 ± 0.003 mm^2^, 0.092 ± 0.003 mm^2^, and 0.093 ± 0.004 mm^2^, respectively), day 4 (0.244 ± 0.014 mm^2^, 0.262 ± 0.014 mm^2^, and 0.273 ± 0.016 mm^2^, respectively) or day 6 (0.573 ± 0.019 mm^2^, 0.599 ± 0.018 mm^2^ and 0.591 ± 0.020 mm^2^, respectively; mean ± SEM) of differentiation, despite the markedly reduced *Blimp1*-Venus expression observed in the *Fancm*^−/−^ clones (**Fig. 4a, b**; p-values and effect sizes reported in **Supplementary Table S10**). Furthermore, flow cytometric assessment of the embryoid bodies confirmed there was a substantial reduction in the BV+ PGCLCs in the *Fancm*^−/−^ compared to the founder or *Fancm*^+/+^ cells at day 2 (12.04 ± 1.37%, 9.07 ± 0.46%, and 1.28 ± 0.24%, respectively), day 4 (13.83 ± 1.14%, 12.31 ± 0.59%, and 3.30 ± 0.56%, respectively), and day 6 (9.50 ± 0.85%, 8.71 ± 0.34%, and 2.08 ± 0.33%, respectively; mean ± SEM) (**Fig. 4c, d**; p-values and effect sizes reported in **Supplementary Table S11**).

**Fig. 4.**
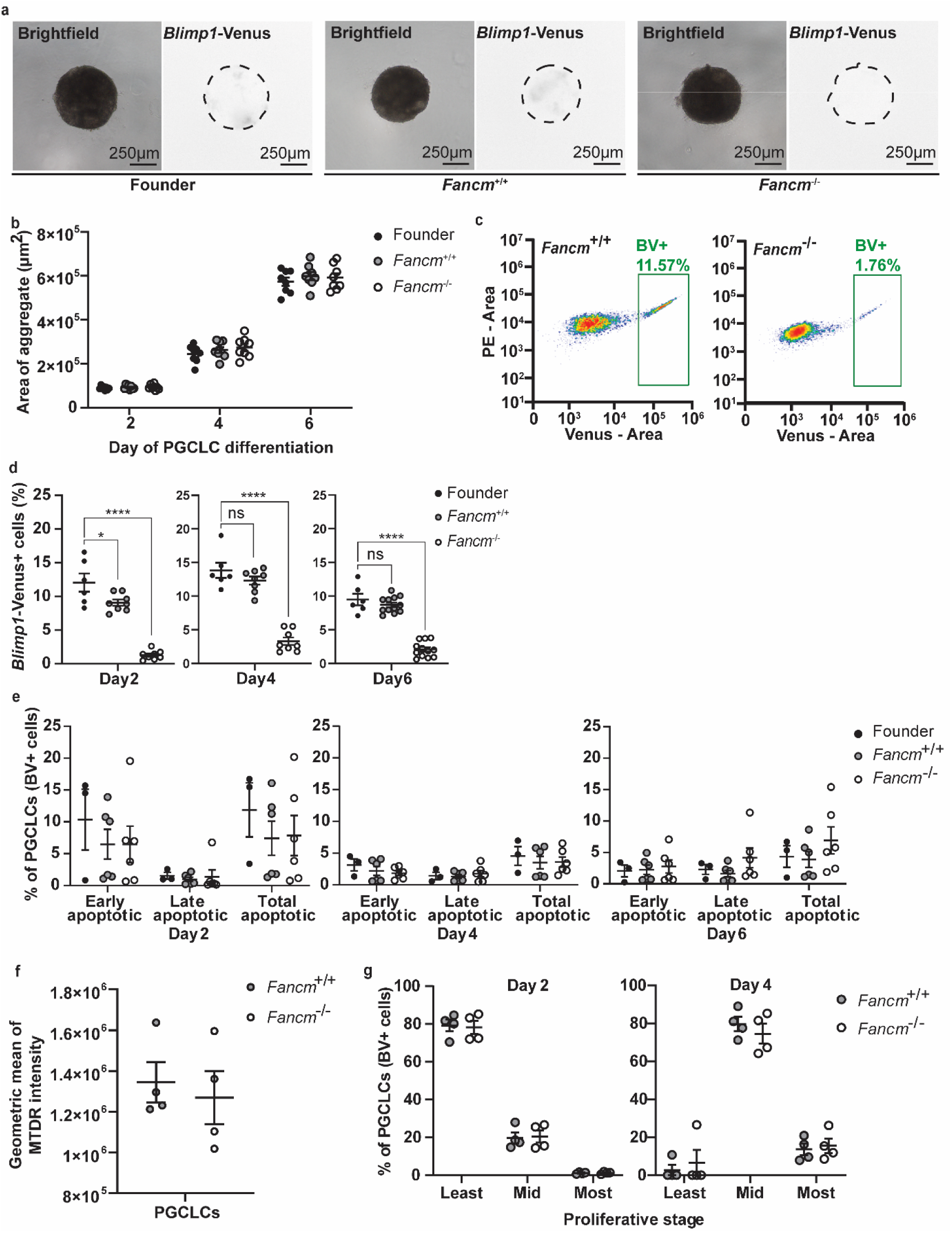
FANCM deficiency compromises germline establishment. (a) Representative brightfield and fluorescent images of founder, *Fancm*^+/+^, and *Fancm*^−/−^ embryoid bodies on day 4 of PGCLC culture. Scale bars = 250 μm. (b) Growth of embryoid bodies measured by quantitative 2D area (μm^2^) of embryoid bodies on day 2, 4, and 6 of PGCLC differentiation. (c) Representative scatter plots and gating for flow cytometric analyses of disassociated embryoid bodies derived from *Fancm*^+/+^ or *Fancm*^−/−^ cell lines, respectively, on day 4 of PGCLC differentiation. PGCLCs are distinguished by expression of the *Blimp1*-Venus transgene (BV+), indicated by rectangular gates. (d) Quantitation of acquisition of BV expression at day 2, 4, and 6 of PGCLC differentiation. * p < 0.05; **p < 0.01; *** p < 0.001; ****p < 0.0001; ns = not significant. (e) Apoptotic profile of BV+ founder, *Fancm*^+/+^ clones, and *Fancm*^−/−^ clones on day 2, 4, and 6 of PGCLC differentiation using Annexin V and propidium iodide (PI) labelling representative of early (Annexin V +, PI -), late (Annexin V + and PI +) and total (Annexin V + and PI −/+) apoptotic cell fractions. (f) Mitochondrial membrane potential of *Fancm*^+/+^ and *Fancm*^−/−^ PGCLCs evaluated by MitoTracker Deep Red (MTDR) staining on day 4 of PGCLC differentiation. (g) Quantitation of analyses of BV+ PGCLC proliferation via CellTrace dye dilution assay, grouped by least-, mid-, and most-proliferative generations.

This reduction was so pronounced that insufficient cells could be harvested for high quality mRNA extraction for qRT-PCR. In contrast, *Fancm*^+/+^ PGCLCs maintained lineage-specific marker expression consistent with the founder line across all time points, ruling out potential off-target effects of CRISPR-Cas9 editing (**Supplementary Fig. S1**; p-values and effect sizes reported in **Supplementary Table S12**).

Given the FA pathway’s role in DNA repair [73], and its relevance in apoptosis and synthetic lethality [74–77], apoptosis was assessed as a potential cause of reduced PGCLC numbers. Early apoptotic markers were highly expressed across all genotypes at day 2 but declined by day 4 and day 6. Overall apoptosis levels were consistent, with no statistically significant differences observed between the founder, Fancm^+/+^ or Fancm^−/−^ PGCLCs at any timepoint (day 2, 4, or 6; **Fig. 4e**; p-values and effect sizes reported in **Supplementary Table S7**).

Next mitochondrial function was investigated, given its importance for PGC survival and function [78,79]. The MitoTracker Deep Red (MTDR) assay was employed to evaluate mitochondrial membrane potential, which serves as a proxy for mitochondrial activity and mass. MTDR staining revealed no differences in mitochondrial membrane potential between *Fancm*^+/+^ and *Fancm*^−/−^ cells at the mESC (**Supplementary Fig. S2a**; p = 0.667, unpaired t-test; *d* = 0.079, very small effect), EpiLC (**Supplementary Fig. S2b**; p = 0.345, unpaired t-test; *d* = −0.561, moderate effect), or PGCLC stage (**Fig. 4f**; p = 0.663, unpaired t-test; *d* = 0.322, small-to-moderate effect).

Cell proliferation was assessed by labelling EpiLCs with a permanent CellTrace dye 30 minutes prior to the start of the six-day PGCLC differentiation protocol. With each cell division, daughter cells inherit approximately half the dye and combining this readout with BV reporter expression enabled cell division tracking within the PGCLCs. Undivided cells or those that were in generation 1-3 were defined as least-proliferative, generation 4-6 as mid-proliferative, and 7-8 as most-proliferative stages (**Supplementary Fig. S3**). As expected, day 2 PGCLCs were largely in the least-proliferative stage and had progressed to the mid-proliferative stage by day 4 (**Fig. 4g**; p-values and effect sizes reported in **Supplementary Table S13**). No proliferation differences were detected between genotypes, despite there being markedly reduced numbers in the *Fancm*^−/−^. These results demonstrate that loss of FANCM results in a severe reduction of PGCLCs, demonstrating that this *in vitro* model system phenocopies *in vivo* mouse models of FANCM loss during germline specification [32,33,80].

## Discussion

POI is a genetically heterogeneous disorder, with increasing evidence implicating genes involved in DNA damage response and repair. Recent reports have identified heterozygous and biallelic pathogenic variants in *FANCM* among individuals with non-syndromic POI (**Supplementary Table S1**), suggesting a potential role for FANCM-mediated genome maintenance during the female reproductive lifespan. However, the developmental stage at which FANCM dysfunction exerts its effects on the germline remains incompletely defined. In this study, we used an *in vitro* PGCLCs differentiation system to assess whether FANCM loss impacts early germline establishment, providing a framework to functionally evaluate POI-associated genetic variants.

This system offers significant advantages over traditional embryo- or foetus-based approaches, most notably in terms of experimental accessibility and temporal resolution. The induction of PGCLCs enables the detailed investigation of early germline specification and lineage commitment under conditions that are readily monitored and manipulated. In contrast to the *in vivo* embryo, where PGCs arise in limited numbers and are located deep within embryonic tissues, PGCLCs can be generated in scalable quantities, synchronized in developmental timing, and sampled at defined intervals.

Previous studies have demonstrated that FANCM deficiency results in reduced fertility and germ cell depletion in a strain-dependent manner (**Supplementary Table S2**). During early foetal development, FANCM-null embryos exhibit a marked reduction in PGC numbers within the foetal gonads at E11.5 to E12.5 [32,33]. This loss is attributed to defective PGC proliferation, rather than increased apoptosis [32,33]. Observed proliferation defects stem from compromised replication-associated DNA repair [32,33,81], rather than from defects in repairing classic interstrand crosslinks. Due to their highly proliferative nature, PGCs are particularly vulnerable to this form of DNA replication stress [32,81,82]. Partial rescue of germ cell numbers in FANCM-null males is achieved by reduction of ATM or loss of TP53 or CDN1A (P21) and in FANCM-null female mice via loss of CHK2 [32,82]. These results indicate that these checkpoint proteins mediate the PGC response to replication stress *in vivo* [32,82].

Consistent with *in vivo* findings, herein *in vitro* analyses show that loss of FANCM does not increase apoptosis in mESCs, EpiLCs, or PGCLCs. Instead, it leads to an almost complete failure of differentiation into the PGCLC state. However, the PGCLC system is comprised of mixed cell populations, and it therefore remains unclear whether this phenotype reflects the requirement of FANCM within the PGCLCs themselves, or indirect effects from surrounding somatic cells in culture. Importantly, the data presented herein suggests that the effects of FANCM loss could occur earlier in germline development than previously recognized. Because PGCLCs correspond to pre-migratory or early migratory PGCs (E7.25-E8.5 *in vivo*), these results imply that FANCM function is critical for the proper specification of early germline progenitors, prior to their colonization of the foetal gonads.

Collectively, these findings reinforce the established role of FANCM in germ cell proliferation while extending its significance to the initial stages of germline specification. The PGCLC model confirms a developmental bottleneck during PGCLC formation that results in a reduced founding germ cell pool. This depletion likely restricts primordial follicle assembly during foetal development, providing a mechanistic link to the pathogenesis of POI.

This study is limited by the inherent genomic instability and aneuploidy of the mESC lines employed, which may influence cellular behaviour and experimental outcomes. However, control (*Fancm*^+/+^) lines carrying identical aneuploidies were used to mitigate confounding effects attributable to chromosomal abnormalities. Moreover, while *in vitro* culture systems lack the physiological complexity and integrative dynamics of whole-organism contexts, the consistency of findings presented herein with results obtained from *in vivo* mouse models supports the robustness and biological relevance of the observed effects. Future studies incorporating targeted validation of replication stress and checkpoint pathways, as well as transcriptomic or single cell approaches, will be required to resolve the cellular basis of the observed phenotype.

Despite these limitations, the PGCLC model contributes a practical refinement of germline research, reducing the need for extensive animal use and complex embryo dissections, while improving experimental reproducibility and scalability. The ability to generate germline-competent cells entirely *in vitro* facilitates cross-species comparisons and lays the groundwork for investigating mechanisms of germline development. PGCLCs provide a powerful and flexible system for elucidating the molecular framework of germline specification and development. The accessibility, controllability, and compatibility with modern molecular and omics-driven tools opens doors for advancing the understanding of early germline development in the mouse and beyond. Looking forward, combining this approach with *in vitro* ovarian organoid systems could enable high-throughput screening of genetic variants linked to premature reproductive aging and other fertility disorders, offering a promising new avenue for research in reproductive health and disease modelling.

## Acknowledgements

We gratefully acknowledge Professor Mitinori Saitou, MD, PhD (Kyoto University, Kyoto, Japan) for his generous donation of the *Blimp1*-Venus *Stella*-ECFP mouse embryonic stem cell line and Bernard de Massy (Centre national de la recherche scientifique (CNRS), Montpellier, France) for his assistance in facilitating the material transfer. We also extend our sincere thanks to the staff of the Flow Cytometry Core at the Novo Nordisk Foundation Center for Stem Cell Medicine (reNEW) for their expert support and guidance. The reNEW Flow Cytometry Platform is funded by the Novo Nordisk Foundation (NNF21CC0073729). Special appreciation is extended to the HERA Consortium for their support, made possible by funding from the Wellcome Trust (316959/Z/24Z). Additional recognition is accorded to Andy Chan for assistance with low-pass sequencing and karyotype analysis.

## Declarations

### Authors’ contributions

Conceptualization: AVK, AP, ERH, and JAH; Funding acquisition: AVK, AP, ERH, and JAH; Supervision: KRSM, ERH, and JAH; Methodology: AVK, HRS, RS, KRSM, and JAH; Investigation: AVK, HRS, RS, and JAH; Data curation: AVK; Formal analysis: AVK and BRP; Visualization: AVK; Writing - original draft preparation: AVK; Writing - review and editing: All authors.

## Supplementary figures

**Supplementary Fig. S1.**
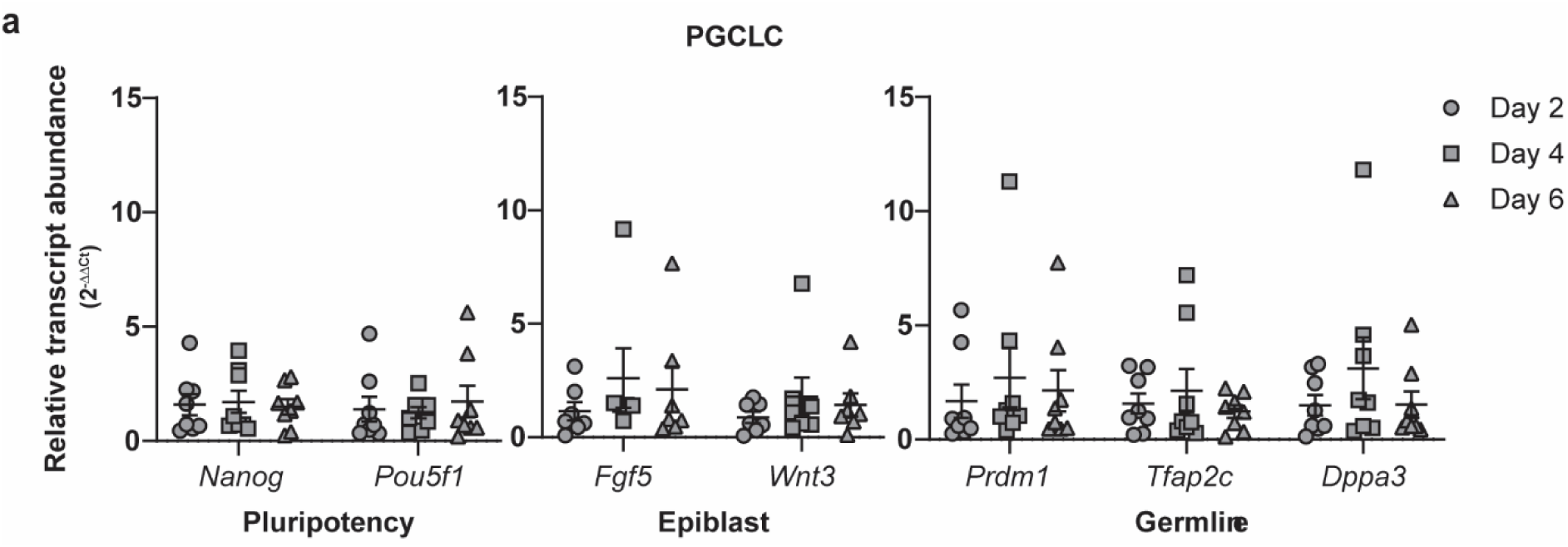
*Fancm*^+/+^ PGCLCs share gene expression profile with euploid founder PGCLCs. (a) Gene expression profile of markers of pluripotent- (*i.e.*, *Nanog*, *Pou5f1*), epiblast- (*i.e.*, *Fgf5*, *Wnt3*), and germline- (*i.e.*, *Prdm1*, *Tfap2c*, and *Dppa3*) lineages of *Fancm*^+/+^ BV+ PGCLCs, normalized to unedited founder BV+ PGCLC expression. Data are presented in log2 scale.

**Supplementary Fig. S2.**
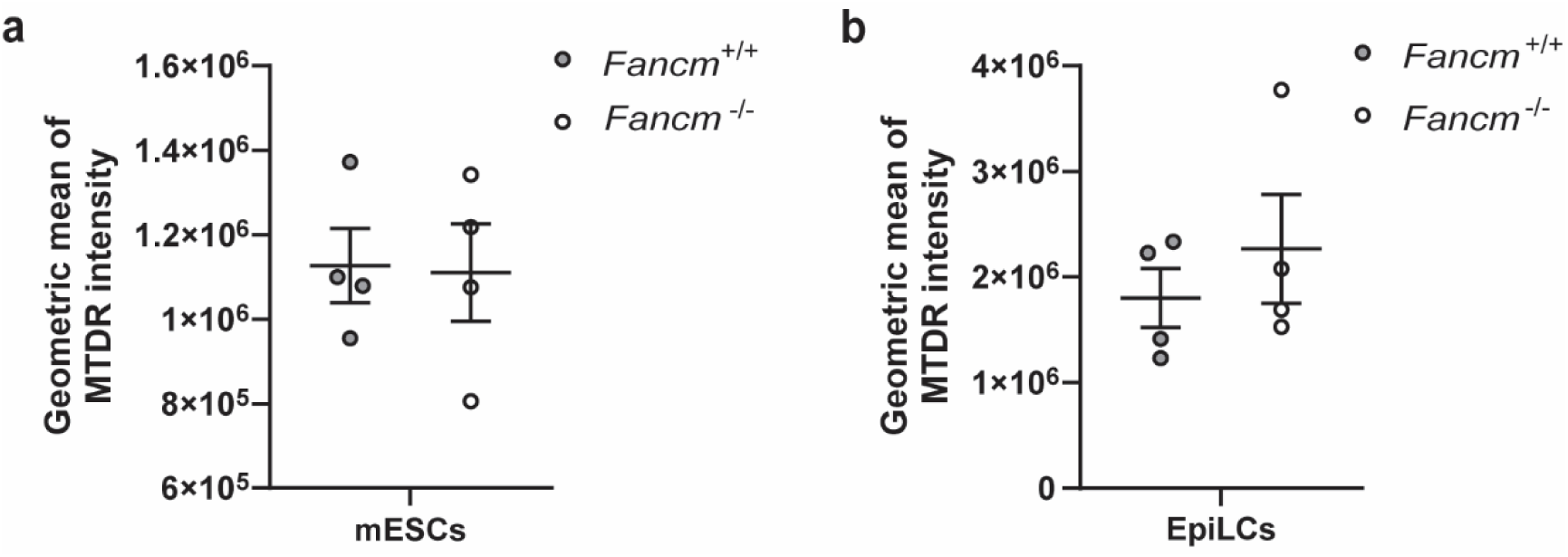
Loss of FANCM does not change mitochondrial membrane potential at pluripotent or epiblast-like stages. (a-b) Mitochondrial membrane potential of *Fancm*^+/+^ and *Fancm*^−/−^mESCs or EpiLCs, respectively, as assessed via MitoTracker Deep Red (MTDR) staining.

**Supplementary Fig. S3.**
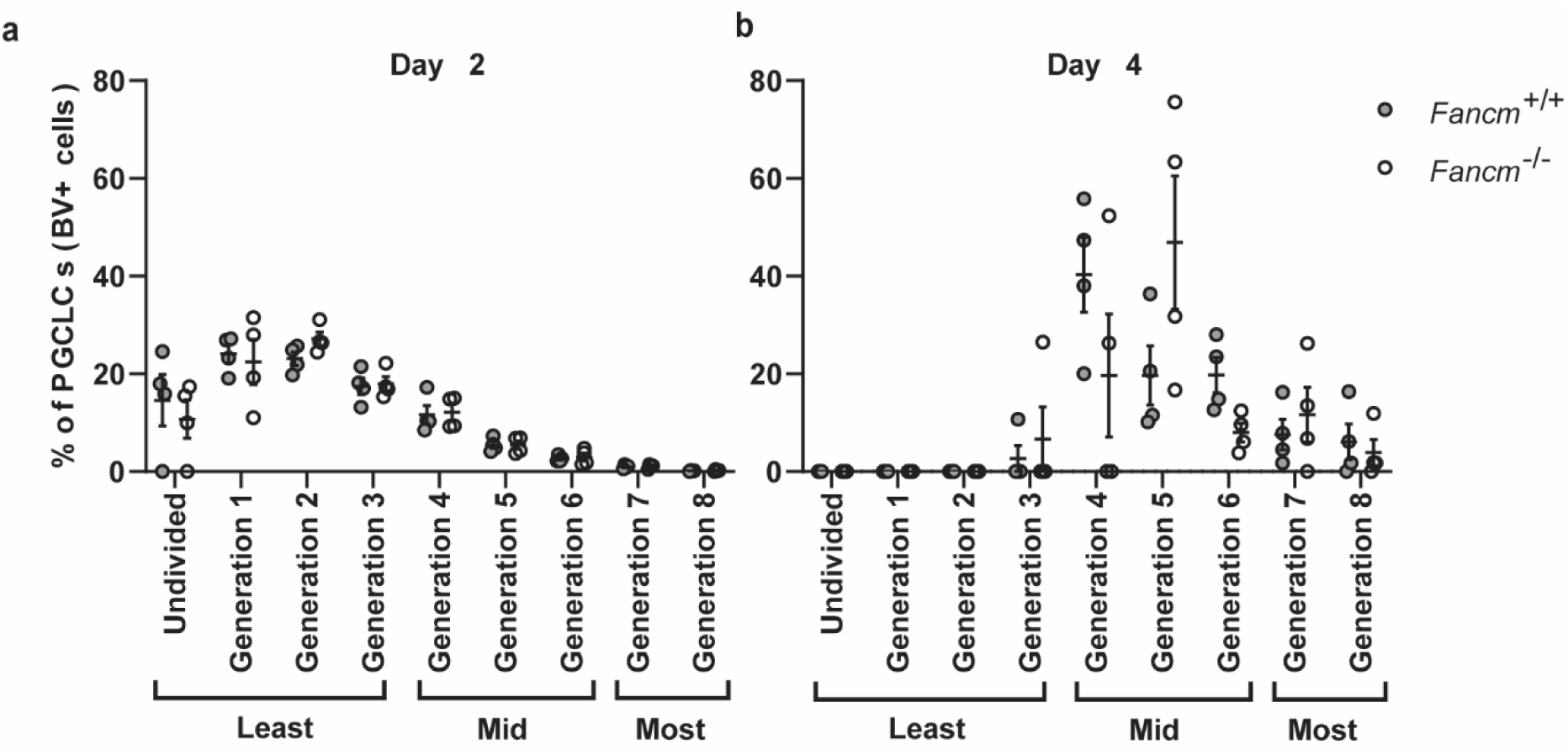
Generational grouping design for CellTrace analysis. (a-b) Representative grouping for least-, mid-, and most-proliferative generations for *Fancm*^+/+^ and *Fancm*^−/−^ PGCLCs at day 2 or 4 of differentiation, respectively, as assessed by CellTrace dye dilution assay.

## Supplementary tables

**Supplementary Table S1.** Summary of literature associating pathogenic *FANCM* variants with non-syndromic premature ovarian insufficiency.

| Variant(s) | Variant type | Variant classification | Allelic status | Variant carriers | Phenotype (age at diagnosis) | Ethnicity | Other potentially causative variants reported | Ref. |
| --- | --- | --- | --- | --- | --- | --- | --- | --- |
| c.362C>T<br>(p.Ala121Val)<br>c.1735C>T<br>(p.Arg579Cys) | Missense<br>Missense | --<br>-- | Heterozygous<sup>a</sup> | 1 | Unspecified | European | Reported as monogenic | [19] |
| c.504G>C<br>(p.Met168Ile) | Missense | Pathogenic | Heterozygous | 1 | PA | -- | Monoallelic missense *ZP1* variant | [24] |
| c.1213C>T<br>(p.Arg405Ter) | Nonsense | Likely pathogenic | Heterozygous | 1 | SA | -- | Reported as monogenic | [22] |
| c.1478G>A<br>(p.Gly519Asp) | Missense | Pathogenic | Heterozygous | 1 | PA | -- | Monoallelic missense variants in *BMP15* & *DCAF17* | [24] |
| c.2215_2216delTG<br>(p.Trp739AlafsTer11) | Frameshift | Pathogenic | Homozygous | 1 | SA (31) | Iranian | Monoallelic nonsense *GNRHR* variant | [17] |
| c.2583_2586<br>delTAAAA:T<br>(p.Lys863IlefsTer12) | Frameshift | Pathogenic | Heterozygous | 1 | Unspecified | European | Reported as monogenic | [19] |
| c.3088C>T<br>(p.Arg1030Ter)<br>c.5791C>T<br>(p.Arg1931Ter) | Nonsense<br>Nonsense | Pathogenic<br>Pathogenic | Heterozygous<sup>a</sup> | 1 | SA (25) | European | Monoallelic missense variants in *AMH* & *MCM8* | [21] |
| c.5101C>T<br>(p.Gln1701Ter) | Nonsense | Pathogenic | Homozygous | 3 | PA | Finnish | Reported as monogenic | [20] |
| c.5101C>T<br>(p.Gln1701Ter) | Nonsense | Pathogenic | Homozygous | 2<sup>b</sup> | SA (22, 24) | Finnish | Reported as monogenic | [18] |
| c.5101C>T<br>(p.Gln1701Ter)<br>c.1636G>A<br>(p.Gly546Ser) | Nonsense<br>Missense | Pathogenic<br>Pathogenic | Heterozygous<sup>a</sup> | 1 | SA (30) | -- | Reported as monogenic | [16] |
| c.5101C>T<br>(p.Gln1701Ter)<br>c.1528G>A<br>(p.Gly510Ser) | Nonsense<br>Missense | Pathogenic<br>VUS | Heterozygous<sup>a</sup> | 1 | SA (18) | European | Monoallelic nonsense *ATM* variant | [20] |
| c.5101C>T<br>(p.Gln1701Ter)<br>c.575A>T<br>(p.Gln192Leu) | Nonsense<br>Missense | Pathogenic<br>Likely pathogenic | Compound heterozygous | 1 | SA (20) | European | Reported as monogenic | [20] |
| c.5653C>T<br>(p.Gln1885Ter) | Nonsense | Likely pathogenic | Heterozygous | 1 | SA | -- | Reported as monogenic | [22] |
| c.5791C>T<br>(p.Arg1931Ter) | Nonsense | Pathogenic | Heterozygous | 1 | PA | -- | Reported as monogenic | [22] |
| c.5791C>T<br>(p.Arg1931Ter) | Nonsense | Pathogenic | Heterozygous | 1 | SA | European | Monoallelic missense *TWNK* variant | [20] |

<sup>a</sup> Two heterozygous variants; cis/trans configuration not reported
<sup>b</sup> Sisters of a consanguineous pedigree

**Supplementary Table S2.** Summary of *in vivo* reproductive consequences of previously published FANCM mouse models.

| Strain designation | Genomic modification | Protein consequence | Strain background | Primordial germ cells | Reproductive consequence | Ref. |
| --- | --- | --- | --- | --- | --- | --- |
| *Fancm*<sup>C4</sup> | Point mutation in exon 1 (c.T524C) | p.C142A within highly conserved DEXDc domain of this DEAD-like helicase superfamily region | C57BL/6J; subsequently backcrossed to C3HeB/FeJ | Reduced germ cell number (at ~E12.5) due to reduced proliferation (not apoptosis) | Smaller litters<br>Female<br>Reduced primordial follicles<br>Reduced germ cells at birth<br>Male<br>Reduced germ cell number<br>Mosaic tubules<br>Spermatocyte<br>DBS repair defect | [32] |
| *Fancm*<sup>Δ2</sup> | Deletion of exon 2 via Cre-lox, introducing frameshift and premature stop codon in exon 3 | Knockout | C57BL/6; subsequently backcrossed to FVB | Reduced germ cell number<br>Reduced germ cell proliferation | Smaller litters<br>Fewer litters<br>Underrepresentation of female pups<br>Female<br>Reduced follicles<br>Male<br>Reduced testis size<br>Reduced germ cell number<br>Mosaic tubules<br>Increased meiotic crossovers | [80,33] |
| *Fancm*<sup>em1/Jcs</sup> | 7 bp frameshift deletion in exon 1 | Knockout | C57BL/6J | -- | Smaller litters<br>Underrepresentation of female pups<br>Male<br>Increased genome instability (exacerbated by concurrent loss of P21) | [82,83] |
| *Fancm*<sup>ΔC</sup> | Frameshift c.1914_1932del mimicking clinically relevant human *FANCM* c.1946_1958del, p.P648Lfs*16 | Truncated protein of 645 aa (p. R638Rfs*8) | C57BL/6 × DBA/2J hybrid; subsequently backcrossed to C57BL/6 | -- | Smaller litters<br>Fewer litters<br>Male<br>Reduced testis size<br>Mosaic tubules<br>Reduced sperm motility<br>Increased sperm morphological abnormalities | [84] |

E: Embryonic day
DSB: Double-strand break

**Supplementary Table S3.**
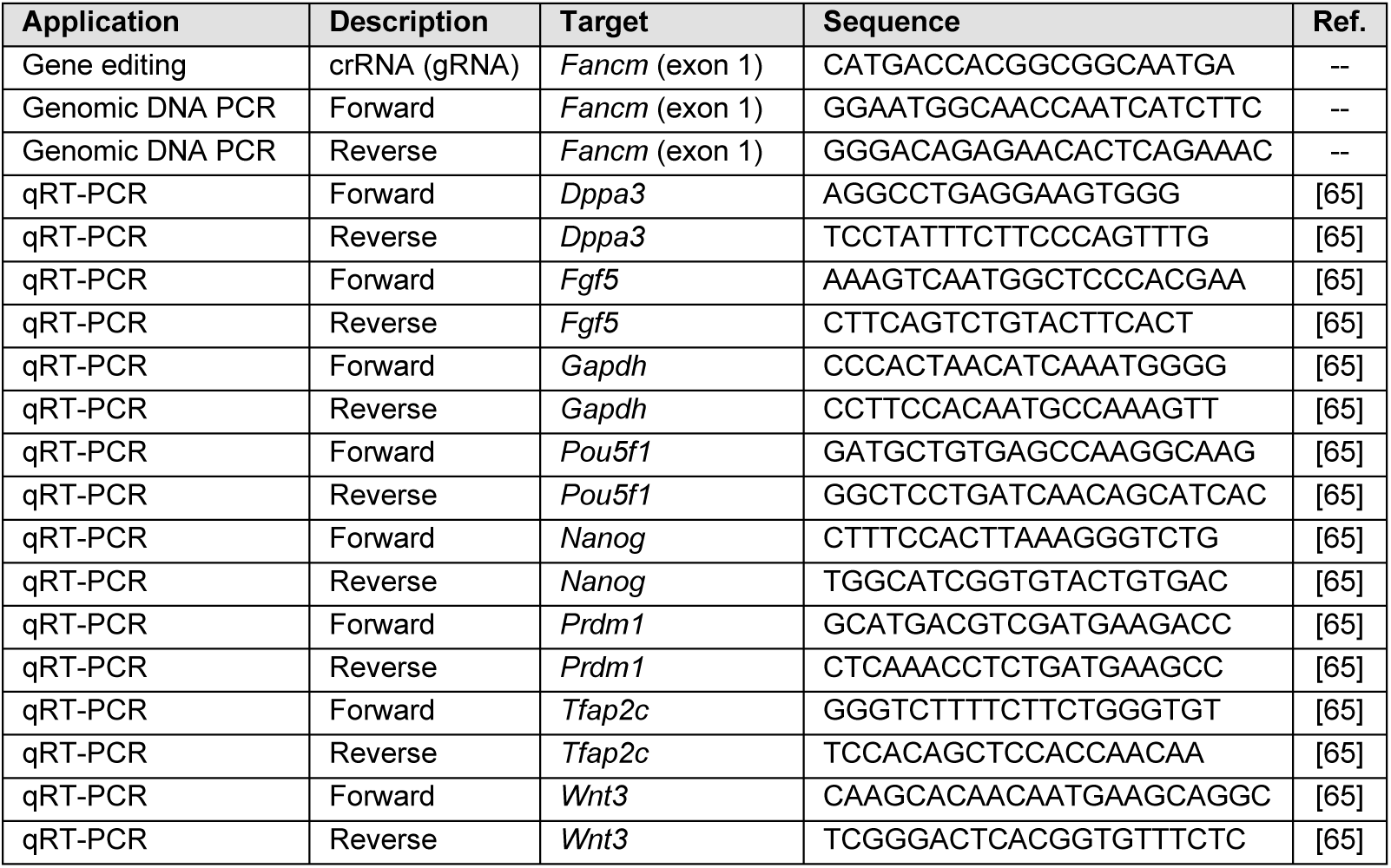
Oligonucleotides used in experimentation.

**Supplementary Table S4.** Gene consequences of CRISPR-Cas9 editing of mouse *Fancm* exon 1.

| Clone | Genotype designation | Deletion (bp) | Insertion (bp) | Change location (nt #) | Classification | Gene annotation (GRCm39:CM001005.3; ENSMUSG00000055884) |
| --- | --- | --- | --- | --- | --- | --- |
| H18 (P21+4+5) | Founder | 0 | 0 | -- | Wildtype | -- |
| H18 (P21+5+2) | Founder | 0 | 0 | -- | Wildtype | -- |
| H18 (P21+3+6) | Founder | 0 | 0 | -- | Wildtype | -- |
| H18 (P21+3+3) | Founder | 0 | 0 | -- | Wildtype | -- |
| Clone E | *Fancm*<sup>+/+</sup> | 0 | 0 | -- | Wildtype | -- |
| Clone G | *Fancm*<sup>+/+</sup> | 0 | 0 | -- | Wildtype | -- |
| Clone I | *Fancm*<sup>+/+</sup> | 0 | 0 | -- | Wildtype | -- |
| Clone Q | *Fancm*<sup>+/+</sup> | 0 | 0 | -- | Wildtype | -- |
| Clone J | *Fancm*<sup>-/-</sup> | 1 | 0 | 324 | Frameshift | *Fancm*:g.324del |
| Clone L | *Fancm*<sup>-/-</sup> | 4 | 0 | 321-324 | Frameshift | *Fancm*:g.321_324del |
| Clone M | *Fancm*<sup>-/-</sup> | 155 | 0 | 173-327 | Frameshift | *Fancm*:g.173-327del |
| Clone W | *Fancm*<sup>-/-</sup> | 111 | 0 | 226-336 | Frameshift | *Fancm*:g.226-336del |

**Supplementary Table S5.** Protein consequences of CRISPR-Cas9 editing of mouse *Fancm* exon 1.

| Clone | Genotype designation | Presence on Western blot | Premature stop codon location (AA #) | 1st exon length (AA) | Total protein length (AA) | Classification | Protein annotation (UniProtKB: Q8BGE5) |
| --- | --- | --- | --- | --- | --- | --- | --- |
| H18 (P21+4+5) | Founder | + | - | 572 | 2,021 | Wildtype | - |
| H18 (P21+5+2) | Founder | + | - | 572 | 2,021 | Wildtype | - |
| H18 (P21+3+6) | Founder | + | - | 572 | 2,021 | Wildtype | - |
| H18 (P21+3+3) | Founder | + | - | 572 | 2,021 | Wildtype | - |
| Clone E | *Fancm*<sup>+/+</sup> | + | - | 572 | 2,021 | Wildtype | - |
| Clone G | *Fancm*<sup>+/+</sup> | + | - | 572 | 2,021 | Wildtype | - |
| Clone I | *Fancm*<sup>+/+</sup> | + | - | 572 | 2,021 | Wildtype | - |
| Clone Q | *Fancm*<sup>+/+</sup> | + | - | 572 | 2,021 | Wildtype | - |
| Clone J | *Fancm*<sup>-/-</sup> | - | 135 | 134 | 134 | Truncation | FANCM:p.Ile108Metfs*27 |
| Clone L | *Fancm*<sup>-/-</sup> | - | 144 | 143 | 143 | Truncation | FANCM:p.Phe107Leufs*37 |
| Clone M | *Fancm*<sup>-/-</sup> | - | 63 | 62 | 62 | Truncation | FANCM:p.Pro58Argfs*5 |
| Clone W | *Fancm*<sup>-/-</sup> | - | 77 | 76 | 76 | Truncation | FANCM:p.Thr75Profs*2 |

**Supplementary Table S6.**
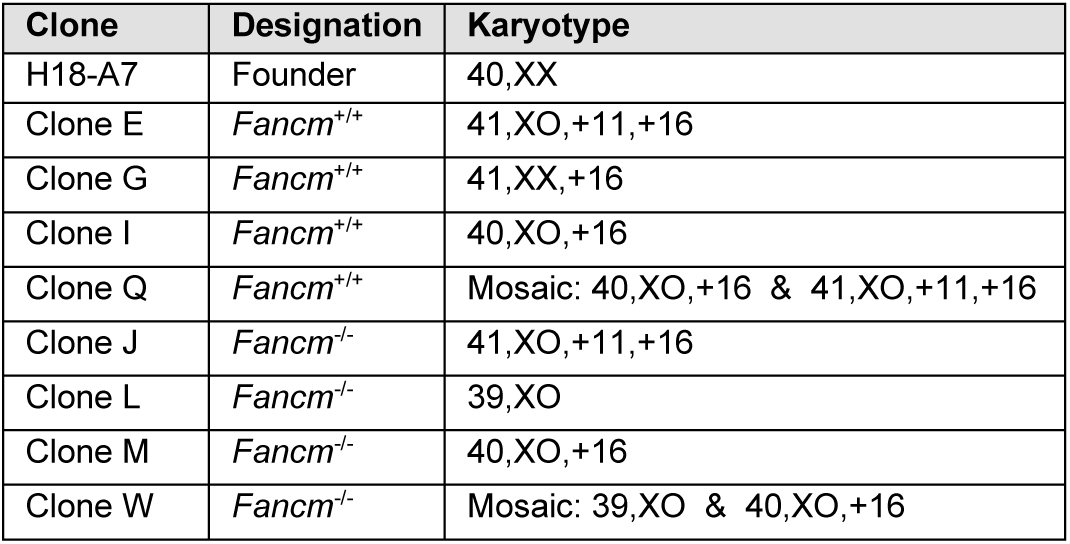
Karyotypes of CRISPR-Cas9 clones.

| Clone | Designation | Karyotype |
| --- | --- | --- |
| H18-A7 | Founder | 40,XX |
| Clone E | *Fancm*<sup>+/+</sup> | 41,XO,+11,+16 |
| Clone G | *Fancm*<sup>+/+</sup> | 41,XX,+16 |
| Clone I | *Fancm*<sup>+/+</sup> | 40,XO,+16 |
| Clone Q | *Fancm*<sup>+/+</sup> | Mosaic: 40,XO,+16 & 41,XO,+11,+16 |
| Clone J | *Fancm*<sup>-/-</sup> | 41,XO,+11,+16 |
| Clone L | *Fancm*<sup>-/-</sup> | 39,XO |
| Clone M | *Fancm*<sup>-/-</sup> | 40,XO,+16 |
| Clone W | *Fancm*<sup>-/-</sup> | Mosaic: 39,XO & 40,XO,+16 |

**Supplementary Table S7.**
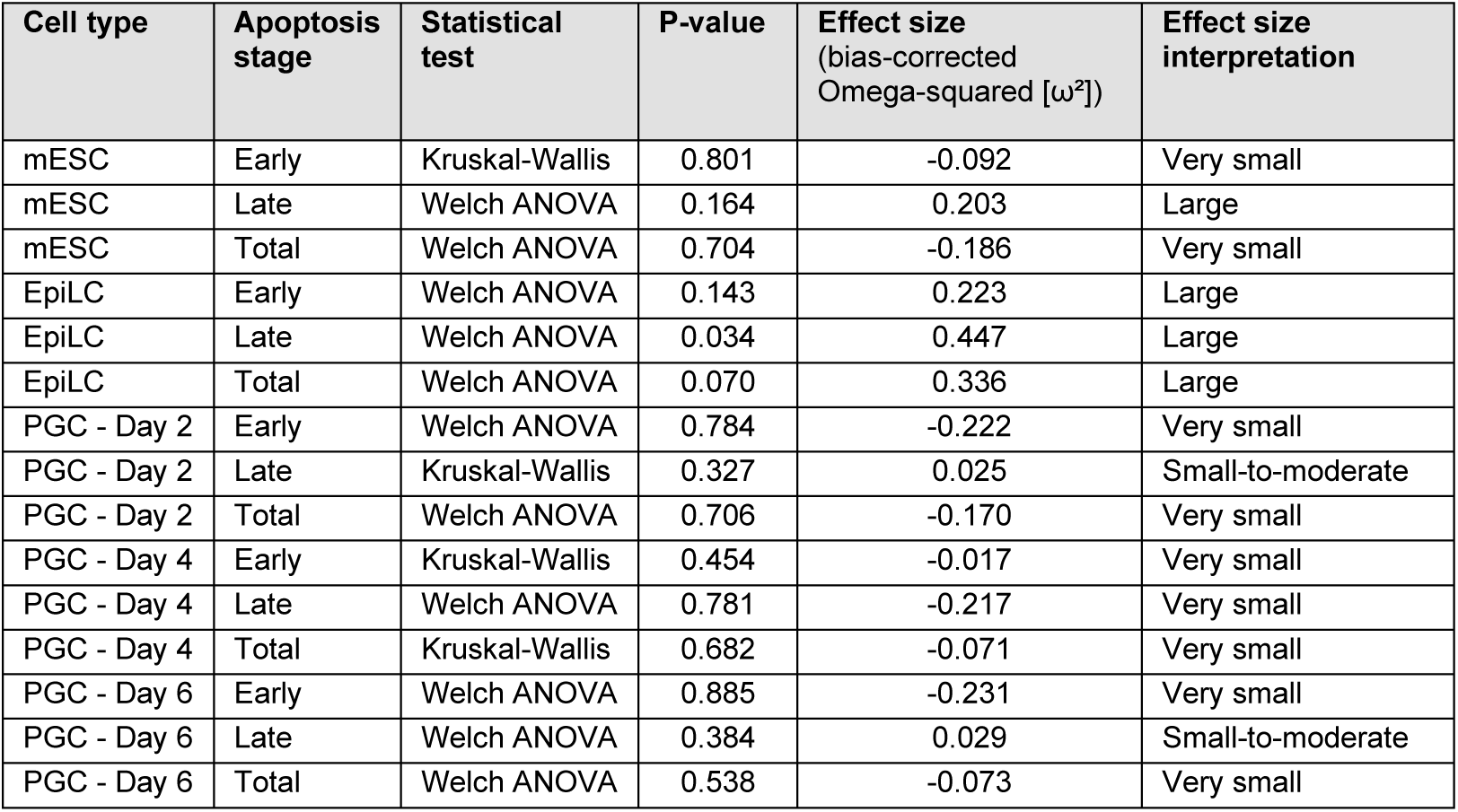
Statistical analyses for apoptosis assays (comparators = Founder, *Fancm*^+/+^, and *Fancm*^−/−^ cell lines)

**Supplementary Table S8.** Post-hoc analysis for significant EpiLC late-stage apoptosis (comparators = Founder, *Fancm*^+/+^, and *Fancm*^−/−^ cell lines) via Games-Howell’s multiple comparisons test

| Comparison | P-value | Effect size (Cohen's *d*) | Effect size interpretation |
| --- | --- | --- | --- |
| Founder vs. *Fancm*<sup>+/+</sup> | 0.101 | -1.673 | Very large |
| Founder vs. *Fancm*<sup>-/-</sup> | 0.089 | -2.084 | Very large |
| *Fancm*<sup>+/+</sup> vs. *Fancm*<sup>-/-</sup> | 0.327 | -0.390 | Small-to-moderate |

**Supplementary Table S9.**
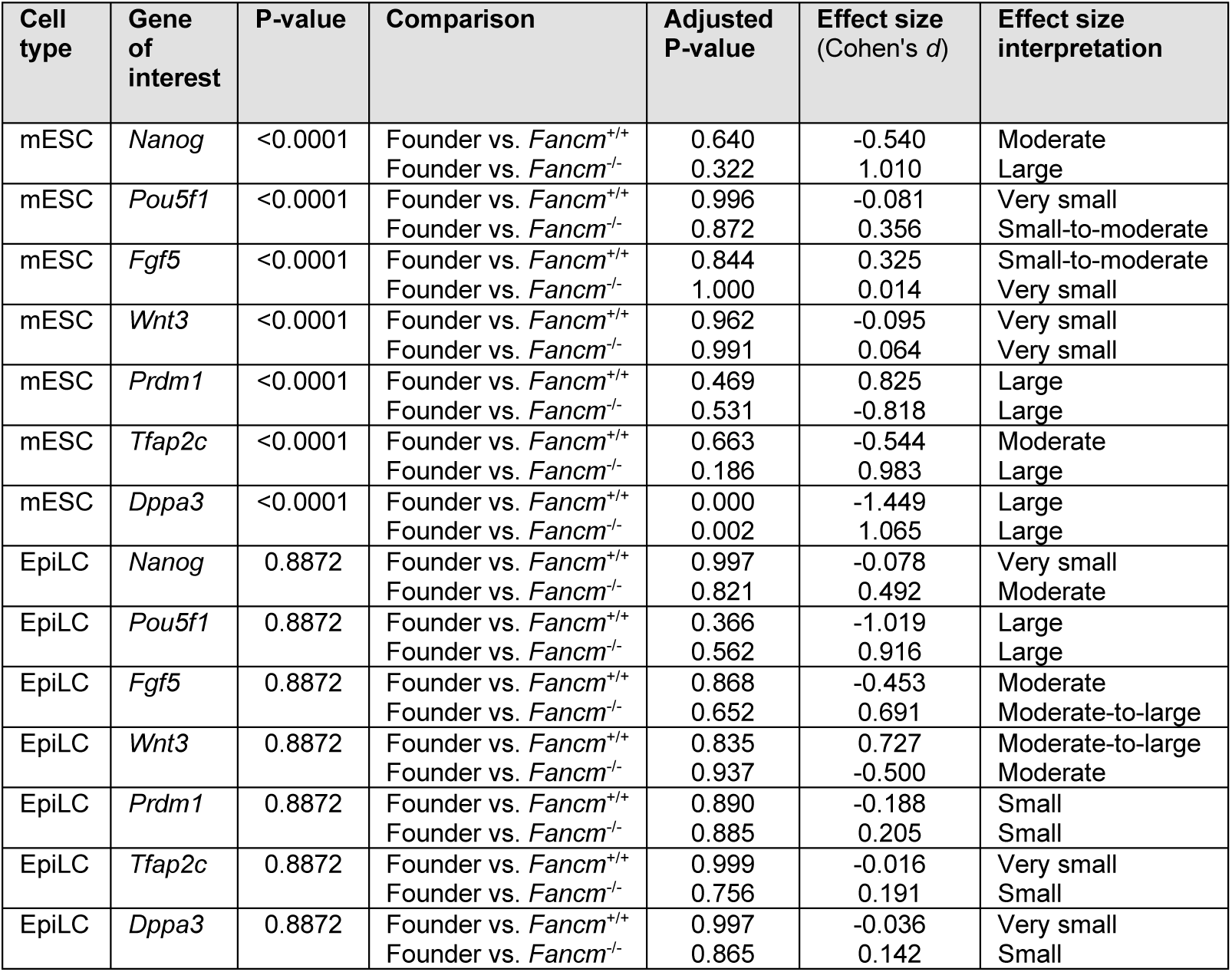
Statistical analyses for qRT-PCR results for mESCs and EpiLCs via 2-way ANOVA; multiple comparisons corrected by statistical hypothesis testing via Dunnett’s multiple comparisons test.

**Supplementary Table S10.** Statistical analyses related to embryoid body size via ordinary one-way ANOVA.

| Day of PGCLC differentiation | P-value | Effect size (bias-corrected Omega-squared [$\omega^2$]) | Effect size interpretation |
| --- | --- | --- | --- |
| Day 2 | 0.683 | -0.054 | Very small |
| Day 4 | 0.391 | -0.001 | Very small |
| Day 6 | 0.620 | -0.044 | Very small |

**Supplementary Table S11.** Statistical analyses for BV+ cell fraction of embryoid bodies via ordinary one-way ANOVA and Tukey’s multiple comparisons test post-hoc test.

| Day of PGCLC differentiation | P-value | Comparison | P-value | Effect size (Cohen's *d*) | Effect size interpretation |
| --- | --- | --- | --- | --- | --- |
| Day 2 | <0.0001 | Founder vs. *Fancm*<sup>+/+</sup> | 0.0258 | 1.257 | Very large |
|  |  | Founder vs. *Fancm*<sup>-/-</sup> | <0.0001 | 4.871 | Very large |
|  |  | *Fancm*<sup>+/+</sup> vs. *Fancm*<sup>-/-</sup> | <0.0001 | 7.871 | Very large |
| Day 4 | <0.0001 | Founder vs. *Fancm*<sup>+/+</sup> | 0.3565 | 0.691 | Moderate-to-large |
|  |  | Founder vs. *Fancm*<sup>-/-</sup> | <0.0001 | 4.856 | Very large |
|  |  | *Fancm*<sup>+/+</sup> vs. *Fancm*<sup>-/-</sup> | <0.0001 | 5.541 | Very large |
| Day 6 | <0.0001 | Founder vs. *Fancm*<sup>+/+</sup> | 0.5044 | 0.482 | Moderate |
|  |  | Founder vs. *Fancm*<sup>-/-</sup> | <0.0001 | 4.598 | Very large |
|  |  | *Fancm*<sup>+/+</sup> vs. *Fancm*<sup>-/-</sup> | <0.0001 | 5.670 | Very large |

**Supplementary Table S12.** Statistical analyses for qRT-PCR results for founder versus *Fancm*^+/+^ PGCLCs via unpaired t-test with multiple comparisons correction (by controlling for False Discovery Rate) via two-stage step-up (Benjamini, Krieger, and Yekutieli)

| Day of PGCLC differentiation | Gene of interest | P-value | Effect size (Cohen's *d*) | Effect size interpretation |
| --- | --- | --- | --- | --- |
| Day 2 | *Nanog* | 0.956 | 0.044 | Very small |
| Day 2 | *Pou5f1* | 0.908 | -0.095 | Very small |
| Day 2 | *Fgf5* | 0.864 | -0.142 | Small |
| Day 2 | *Wnt3* | 0.568 | -0.482 | Moderate |
| Day 2 | *Prdm1* | 0.971 | -0.030 | Very small |
| Day 2 | *Tfap2c* | 0.934 | 0.068 | Very small |
| Day 2 | *Dppa3* | 0.961 | -0.041 | Very small |
| Day 4 | *Nanog* | 0.693 | 0.323 | Small-to-moderate |
| Day 4 | *Pou5f1* | 0.963 | 0.038 | Very small |
| Day 4 | *Fgf5* | 0.543 | 0.531 | Moderate |
| Day 4 | *Wnt3* | 0.850 | 0.154 | Small |
| Day 4 | *Prdm1* | 0.842 | 0.163 | Small |
| Day 4 | *Tfap2c* | 0.933 | 0.069 | Very small |
| Day 4 | *Dppa3* | 0.561 | 0.480 | Moderate |
| Day 2 | *Nanog* | 0.853 | 0.152 | Small |
| Day 2 | *Pou5f1* | 0.981 | 0.019 | Very small |
| Day 2 | *Fgf5* | 0.613 | 0.425 | Moderate |
| Day 2 | *Wnt3* | 0.991 | 0.013 | Small |
| Day 2 | *Prdm1* | 0.713 | 0.301 | Small-to-moderate |
| Day 2 | *Tfap2c* | 0.928 | -0.074 | Very small |
| Day 2 | *Dppa3* | 0.954 | 0.047 | Very small |

**Supplementary Table S13.**
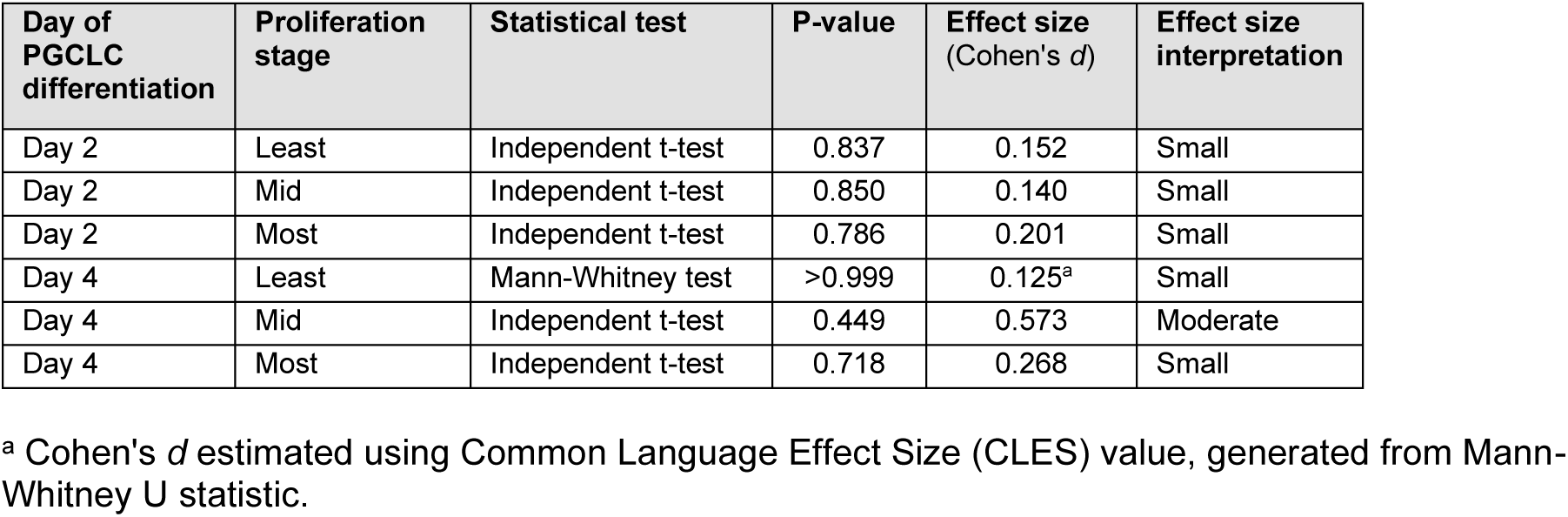
Statistical analyses for PGCLC proliferation assays.

| Day of PGCLC differentiation | Proliferation stage | Statistical test | P-value | Effect size (Cohen's *d*) | Effect size interpretation |
| --- | --- | --- | --- | --- | --- |
| Day 2 | Least | Independent t-test | 0.837 | 0.152 | Small |
| Day 2 | Mid | Independent t-test | 0.850 | 0.140 | Small |
| Day 2 | Most | Independent t-test | 0.786 | 0.201 | Small |
| Day 4 | Least | Mann-Whitney test | >0.999 | 0.125<sup>a</sup> | Small |
| Day 4 | Mid | Independent t-test | 0.449 | 0.573 | Moderate |
| Day 4 | Most | Independent t-test | 0.718 | 0.268 | Small |

<sup>a</sup> Cohen's *d* estimated using Common Language Effect Size (CLES) value, generated from Mann-Whitney U statistic.

